# Heat Hardening of a Larval Amphibian is Dependent on Acclimation Period and Temperature

**DOI:** 10.1101/2022.09.19.508599

**Authors:** Jason Dallas, Robin W. Warne

## Abstract

The thermal tolerance–plasticity trade-off hypothesis states that acclimation to warmer environments increases basal thermal tolerance in ectotherms but reduces plasticity in coping with acute thermal stress characterized as heat hardening. We examined the potential trade-off between basal heat tolerance and hardening plasticity, measured as critical thermal maximum (CT_max_) of a larval amphibian, *Lithobates sylvaticus*, in response to differing acclimation temperatures (15° and 25°C) and periods (3 or 7 days). A hardening treatment applied 2 hours before CT_max_ assays induced pronounced plastic hardening responses in the cool, 15°C treatment after 7 days of acclimation, compared to controls. Warm acclimated larvae at 25°C, by contrast, exhibited minor hardening responses, but significantly increased basal thermal tolerance. These results support the trade-off hypothesis and fill a knowledge gap in larval amphibian thermal plasticity. Elevated environmental temperatures induce acclimation in heat tolerance yet constrains ectotherm capacity to cope with further acute thermal stress.

**Summary Statement:** A larval amphibian follows the trade-off hypothesis such that the group with the highest basal heat tolerance displays the lowest hardening response and vice-versa.

## Introduction

Environmental temperature is one of the most important abiotic drivers of organismal physiology (Angilletta Jr, 2009). As temperatures increase due to climate change, ectotherms will be under greater risk of approaching their upper thermal limit that will lead to shifts in species distributions, altered biological interactions, and reduced activity periods, all of which can result in extinction (Bellard et al., 2012; Blois et al., 2013; Cox et al., 2022; Somero, 2010). Global declines in amphibians have been linked to climate change (e.g., Blaustein et al., 2010; Campbell Grant et al., 2020; Lowe et al., 2021; Rollins-Smith, 2017), highlighting the need for continued research on how they respond to warming and thermal extremes.

As thermal traits generally evolve slowly in herpetofauna (Bodensteiner et al., 2020), phenotypic plasticity is likely a primary response to climate change and increasing thermal stress. Thermal acclimation represents reversible plasticity in basal heat tolerance and develops over days to weeks of chronic exposure to altered environmental temperatures (e.g., Cupp Jr, 1980; Lapwong et al., 2021b; Li et al., 2009; Rohr et al., 2018; Sgro et al., 2010). However, acclimation does not necessarily protect organisms against acute exposure to short-term heat events such as heat waves which are projected to increase in frequency (Seneviratne et al., 2021). The related heat hardening response is another form of thermal plasticity that, by contrast, develops rapidly over minutes to hours of exposure to acute heat stress (Bowler, 2005). Heat hardening is generated by exposing organisms to temperatures near or at their upper thermal limit. While hardening rapidly increases heat tolerance, these increases are transient and disappear within 36 hours (Deery et al., 2021; Maness and Hutchison, 1980; Phillips et al., 2016; Rutledge et al., 1987; but see Moyen et al., 2020), highlighting its role as a short-term protection mechanism. Therefore, plasticity in heat tolerance occurs at two different levels: basal thermal tolerance, measured as the limits of thermal performance curves (Huey and Stevenson, 1979), following an acclimation period, and hardening, which temporarily increases basal thermal tolerance following an acute heat stress.

Under an ideal scenario, both high basal thermal tolerance and hardening would improve ectotherm persistence under climate change. However, there appears to be a physiological limitation such that elevated basal thermal tolerance constrains the capacity of an organism to further increase their heat tolerance. For example, Stillman (2003) found a negative relationship between basal thermal tolerance and acclimation capacity in *Petrolisthes* crab populations across a latitudinal gradient. Building upon this, van Heerwaarden and Kellermann (2020) identified that this negative link was widespread across ectothermic clades and named this pattern the tolerance–plasticity trade-off hypothesis. Heat shock proteins (HSPs) may underlie the trade-off hypothesis because of the central role they play in maintaining homeostasis during extreme temperatures (Feder and Hofmann, 1999; Sørensen et al., 2003) and improving basal thermal tolerance (Bahrndorff et al., 2009; Blair and Glover, 2019; Gao et al., 2014; Krebs and Feder, 1998; but see Easton et al., 1987; Jensen et al., 2010). Because HSPs are energetically expensive to produce and maintain (e.g., Hoekstra and Montooth, 2013), populations from warm environments may be ‘preadapted’ to favor relatively high constitutive HSP expression to elevate basal thermal tolerance but exhibit less flexibility in upregulation following an acute heat shock compared to cool environment populations (Gleason and Burton, 2015). Therefore, under the trade-off hypothesis, hardening may be more useful to species that are less likely to experience chronic heat stress but receive greater benefits in combating acute stress (van Heerwaarden and Kellermann, 2020; but see Sgro et al., 2010). Thus, acute upper thermal limits that are near or pushed near adapted thermal maxima restrict additional plasticity for further increased thermal tolerance through acclimation (Somero, 2010). While a meta-analysis on ectotherms by Barley et al. (2021) provided support for the trade-off hypothesis, there is mixed evidence in larval amphibians (e.g., Menke and Claussen, 1982; Simon et al., 2015; Turriago et al., 2022) suggesting a need for further exploration.

The role of heat hardening in adult (Maness and Hutchison, 1980) and larval amphibians (Sherman and Levitis, 2003; Sørensen et al., 2009) is understudied. We aimed to investigate the trade-off hypothesis by testing how acclimation temperatures (low or high) and duration (short or long acclimation periods) affect interactions between heat hardening and basal thermal tolerance - estimated via critical thermal maximum (CT_max_). These tests were conducted on larval wood frogs, *Lithobates sylvaticus* (LeConte 1825). Because larval anurans display a positive relationship between acclimation temperature and CT_max_ (e.g., Cupp Jr, 1980; Ruthsatz et al., 2022), we predicted that longer acclimation to warmer temperatures would increase basal heat tolerance compared to those acclimated to cooler temperatures. In line with the trade-off hypothesis, we also expected the hardening effect would be most pronounced in larvae with the lowest CT_max_ suggesting greater acute thermal plasticity under these environments.

## Materials and Methods

### Field Collection and Husbandry

Freshly laid (< 36 hours old) wood frog egg masses were collected from wetlands in Jackson Co., IL under an Illinois Department of Natural Resources permit (HSCP 19-03). The egg masses were maintained in 60 L plastic containers with aerated, carbon-filtered water. After hatching, larvae were initially fed autoclaved algal flakes (Bug Bites Spirulina Flakes, Fluval Aquatics, Mansfield, MA, USA), followed by crushed alfalfa pellets at two weeks after hatching. Animals were fed twice weekly, and water was changed weekly. All experimental procedures were approved by the Southern Illinois University Institutional Animal Care and Use Committee (22–008).

### Critical Thermal Maximum Assay

After larvae reached early pro-metamorphic stages, 64 individuals were randomly selected and staged, weighed, and transferred to individual 750 mL plastic containers filled with 600 mL of aged (>24 hours) aerated, carbon-filtered water. Larvae were split (N=32/treatment) into low (15°C ± 0.2) and high (25°C ± 0.3) acclimation temperatures. There were no differences in initial Gosner (1960) stage (range = 27 – 35) or mass (0.25 – 0.55 g) between these groups (P > 0.3). The larvae were further randomly split into four groups (n=8 per group) that differed in acclimation period and heat hardening treatment: 1) 3-day control, 2) 3-day hardened, 3) 7-day control, and 4) 7-day hardened. Larvae were acclimated to low or high temperatures for either three or seven days. On the last day of acclimation, the CT_max_ of control groups was measured. The hardened groups were heated for 10 minutes at 2–4°C below the CT_max_ of control groups, following the protocol of Sherman and Levitis (2003). After this heat hardening treatment, the animals were returned to their acclimation temperatures for 2 hours, after which their CT_ma_x was measured. Sample sizes were reduced to seven for the 7-day hardened low and high temperatures groups, and the 7-day control low temperature group due to mortality.

CT_max_ was measured between 1000 – 1600 hrs to minimize potential diel effects on heat tolerance (Healy and Schulte, 2012; Maness and Hutchison, 1980). Larvae were staged, weighed, and then placed in individual 125 mL flasks filled with 75 mL of aged, aerated, carbon-filtered water and submerged in a hot water bath (Isotemp 220, Fischer Scientific) and given 5 minutes to acclimate prior to beginning the assay. Water temperatures increased 0.6 ± 0.01°C per minute from a starting temperature of 19.9 ± 0.2°C. Beginning at ~34°C, larvae were prodded with a spatula every 30 seconds until they did not respond to the stimulus. At this point, a thermocouple probe (Physitemp BAT-12) was placed in the flask, water temperature was recorded which represented the larval CT_max_. Flasks were then placed in a water bath at room temperature to facilitate larval recovery, and all larvae recovered ≤ 5 minutes. Upon completion of CT_max_ measurements, all larvae were euthanized via snap-freezing in −80°C ethanol.

### Statistical Analyses

We assessed how larval CT_max_ shifted in response to our various treatments using a general linear model. While Gosner stage recorded prior to the CT_max_ measurement was normally distributed, mass was log-transformed to achieve normality, and both were included as covariates in the model. Fixed effects included acclimation period (3 or 7 days), acclimation temperature (low or high), hardening treatment (control or hardened), and their interactions. Post-hoc analyses were conducted using Tukey tests. All analyses were conducted in R Studio v. 2022.02.3 (https://www.Rstudio.com/) and significance values were set as α = 0.05.

## Results

Across all treatments, wood frog larvae displayed a moderate degree of variation in their CT_max_ (range = 35.8° – 39.6°C; Table 1). Two individuals were dropped from analyses due to abnormally low CT_max_ values (≤ 34.9°C) in relation to their group mean. Of the main effects, only acclimation temperature (F_1,49_ = 6.52, P = 0.014) had a significant effect (Table 2) with those in the high acclimation temperature treatment exhibiting greater heat tolerance (Fig. 1). While neither hardening (F_1,49_ = 0.088, P = 0.77) nor acclimation period (F_1,49_ = 2.55, P = 0.12) had significant effects on CT_max_, there was significant hardening by acclimation period (F_1,49_ = 6.11, P = 0.017) and acclimation period by acclimation temperature (F_1,49_ = 18.71, P < 0.0001) interactions. The former was driven by a more pronounced hardening effect for day 7 individuals, while the latter was the outcome of a pronounced increase in CT_max_ among larvae in the high acclimation treatment on day 7 (Fig. 1). Lastly, a significant three-way interaction was found for acclimation period, acclimation temperature, and hardening treatment (F_1,49_ = 4.47, P = 0.040). Larvae in the low acclimation treatment on day 7 showed the largest hardening effect of 0.9°C, which was more than double the hardening effect of any other group (Fig. 1). Larval mass and Gosner stage were unrelated to CT_max_ (P ≥ 0.29).

**Figure 1:**
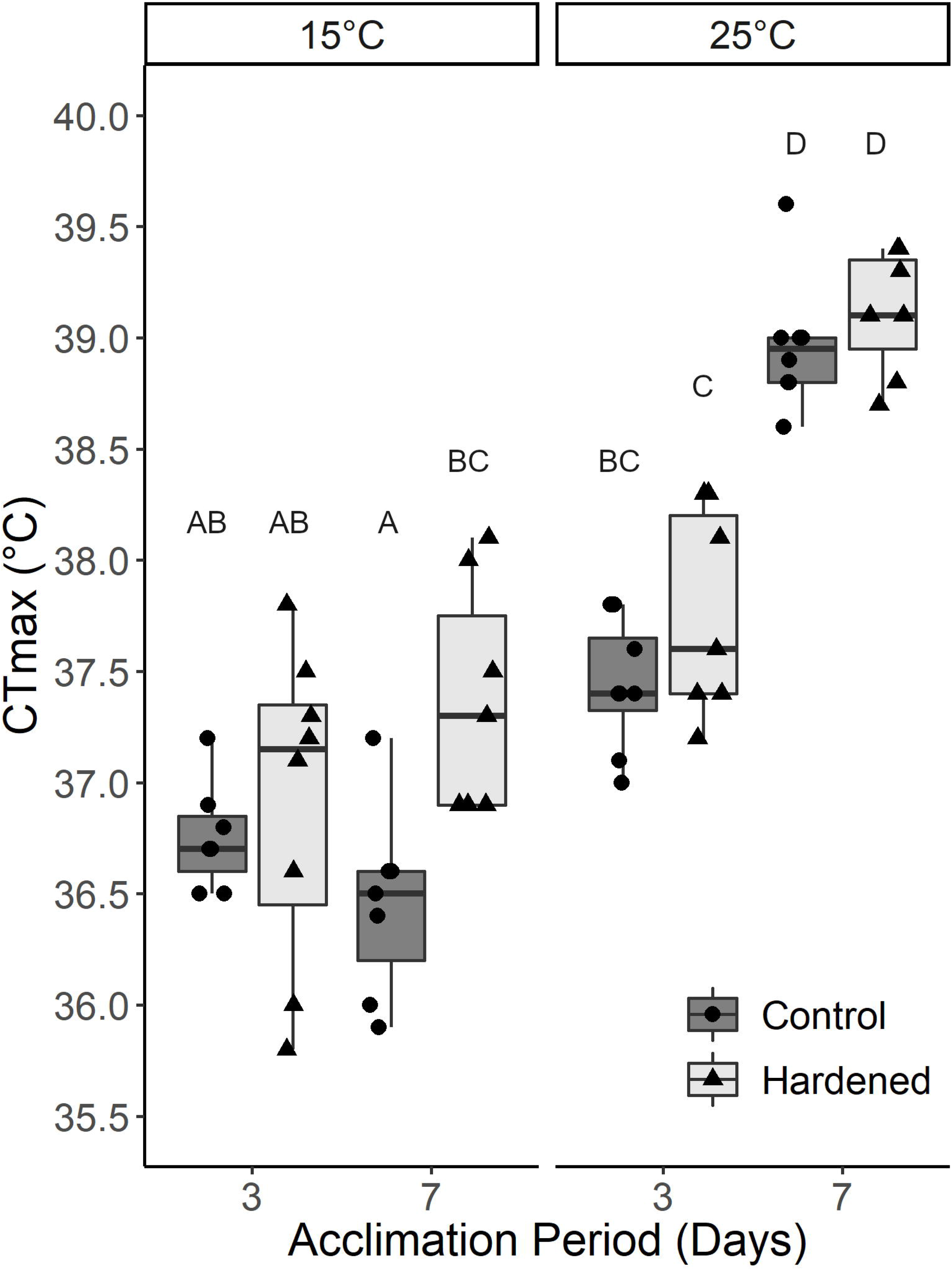
Heat tolerance of larval wood frogs across differing acclimation conditions and hardening. Larval wood frog critical thermal maximum (CT_max_) exposed to two different acclimation temperatures (15° and 25°C), two different acclimation periods (3 and 7 days), and a hardening treatment (control vs. hardened). Points represent individual larvae. Center lines within boxplots represent the median and the boxes denote the interquartile range with whiskers representing 1.5x the upper or lower quartile. Letters indicate significant differences between post hoc pairwise comparisons (Tukey HSD, P < 0.05).

**Table 1:**
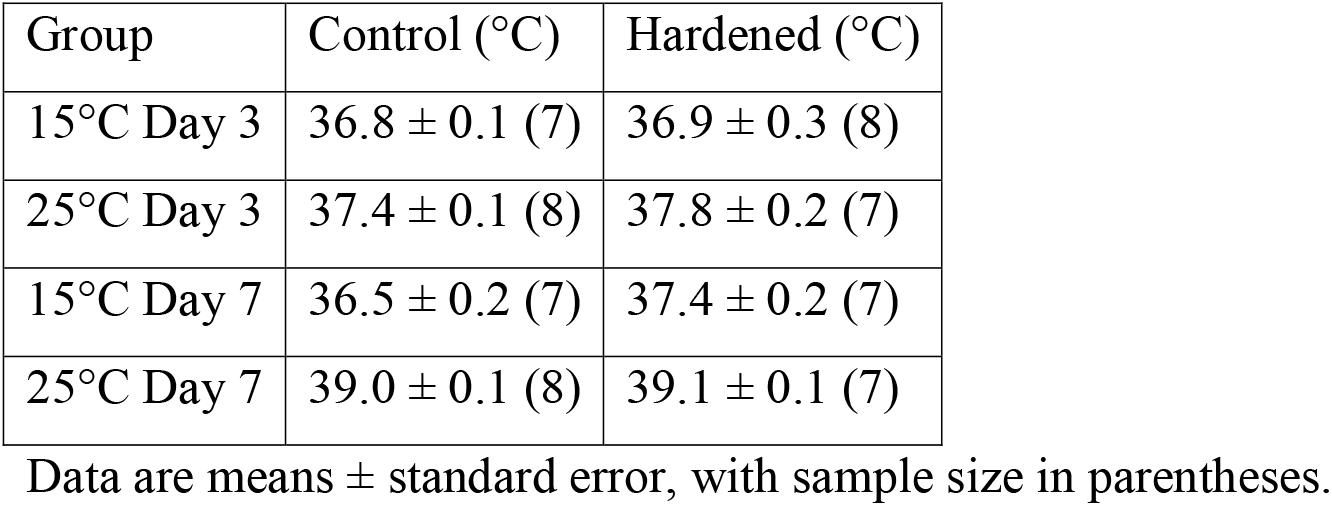
Mean critical thermal maximum (CT_max_) of control and hardened larval wood frogs at different acclimation temperatures and acclimation periods.

| Group | Control (°C) | Hardened (°C) |
| --- | --- | --- |
| 15°C Day 3 | 36.8 ± 0.1 (7) | 36.9 ± 0.3 (8) |
| 25°C Day 3 | 37.4 ± 0.1 (8) | 37.8 ± 0.2 (7) |
| 15°C Day 7 | 36.5 ± 0.2 (7) | 37.4 ± 0.2 (7) |
| 25°C Day 7 | 39.0 ± 0.1 (8) | 39.1 ± 0.1 (7) |

Data are means ± standard error, with sample size in parentheses.

**Table 2:** Effects of body mass, Gosner stage, hardening treatment, acclimation period, and acclimation temperature on larval wood frog critical thermal maximum from a generalized linear model.

| Source of Variation | S. S. | d. f. | F | *P* |
| --- | --- | --- | --- | --- |
| Log-Transformed Body Mass | 0.14 | 1 | 0.73 | 0.40 |
| Gosner Stage | 0.22 | 1 | 1.15 | 0.29 |
| Hardening | 0.017 | 1 | 0.088 | 0.77 |
| Acclimation Period | 0.49 | 1 | 2.55 | 0.12 |
| Acclimation Temperature | 1.25 | 1 | 6.52 | **0.014** |
| Hardening x Acclimation Period | 1.17 | 1 | 6.11 | **0.017** |
| Hardening x Acclimation Temperature | 0.094 | 1 | 0.49 | 0.49 |
| Acclimation Period x Acclimation Temperature | 3.57 | 1 | 18.71 | **< 0.001** |
| Hardening x Acclimation Period x Acclimation Temperature | 0.85 | 1 | 4.47 | **0.040** |
| Residuals | 9.35 | 49 |  |  |

Values in bold indicate significant differences, $p < 0.05$ .

## Discussion

Phenotypic plasticity of heat tolerance provides ectotherms the ability to counter the threat of overheating due to temperature extremes associated with climate change. Heat hardening, a form of thermal plasticity, represents the “first line of defense” against heat stress (Deery et al., 2021) through rapid upregulation of HSPs and/or changes to cellular structure in response to an acute thermal shock that can increase short-term heat tolerance (Bowler, 2005). However, the tolerance–plasticity trade-off hypothesis (van Heerwaarden and Kellermann, 2020) proposes that basal heat tolerance and thermal plasticity are negatively correlated; such that individuals with high CT_max_ have limited hardening (Gilbert and Miles, 2019). While numerous studies have demonstrated that amphibians exhibit plastic basal heat tolerance (e.g., Cupp Jr, 1980; Ruthsatz et al., 2022), hardening remains understudied.

In our study, we found evidence in support of the trade-off hypothesis for larval wood frogs, although the effect was minor (Fig. 1), potentially due to low sample sizes. The group with the lowest mean CT_max_ (36.5°C) had the greatest hardening effect (0.9°C), while the group with the highest mean CT_max_ (39.0°C) had a minimal hardening effect (0.1°C). While the 0.9°C hardening effect was comparable to larval American toads (*Anaxyrus americanus*) and African clawed frogs (*Xenopus laevis*) (Sherman and Levitis, 2003), the remaining groups had a minor hardening response (≤ 0.4°C) that was similar with values for larval bullfrogs (*L. catesbeianus*)(Menke and Claussen, 1982). Additionally, the bullfrogs showed no evidence of the trade-off hypothesis as CT_max_ increased positively with acclimation temperatures while hardening effect was unchanged. Hardening effects in lizard, salamander, and fish species are variable ranging from –0.4°C (*Anolis sagrei*) to 2.1°C (*A. carolinensis*) (Deery et al., 2021; Lapwong et al., 2021a; Maness and Hutchison, 1980; Phillips et al., 2016; Rutledge et al., 1987). In relation to other species, larval wood frogs acclimated to cooler conditions have a relatively strong hardening effect indicating significant plasticity in heat tolerance to improve their tolerance of overheating. This may benefit wood frogs as ephemeral pond breeding species are threatened by climate change (Blaustein et al., 2010) during the larval stage (Enriquez-Urzelai et al., 2019).

We can only speculate on the mechanism that drove the observed results, but we propose that HSPs represent an intriguing answer. This is because they are intimately tied to environmental temperature (Dalvi et al., 2012; Gu et al., 2019; Jin et al., 2019) and basal thermotolerance (Bahrndorff et al., 2009; Blair and Glover, 2019). Warm-tolerant ectotherms often express higher constitutive levels of *hsp70* relative to less-tolerant populations, but that an acute heat-stress results in greater *hsp70* expression in those with lower basal thermal tolerance (Gleason and Burton, 2015; Zatsepina et al., 2000; Zatsepina et al., 2001). Zatsepina et al. (2000) proposed that this provided temperate populations the capability to rapidly and intensely synthesize HSPs after brief exposure to heat shock that was absent in low latitude populations. We propose a similar pattern in the wood frog larvae, such that higher constitutive HSP levels in warm-acclimated larvae provided increased basal heat tolerance compared to cold-acclimated larvae, yet hardened larvae from the latter group greatly upregulated HSP expression following a heat shock enhancing their hardening response. This is in line with *Drosophila* acclimated to cooler temperatures (Bettencourt et al., 1999), which exhibited pronounced hardening plasticity that was absent in the warm-acclimated group. Quantifying constitutive and heat-shocked *hsp70* mRNA of larval liver and gill tissues would offer support to this conclusion. Additionally, many ectotherms appear to have hard upper-limits to thermal tolerance after which their pejus range constraints any further plastic responses (Denny and Dowd, 2012). Thus, the warm acclimated larvae in our study could have approached their physiologically and evolutionarily determined upper limit that constrained any further plastic responses. Future tests are required to understand 1) if there is a degree of plasticity to hard upper limits of thermal acclimation, 2) the cellular and physiological mechanisms underlying these limits, 3) how these mechanisms determine the trade-offs between hardening and acclimation to chronic heat stress, and 4) how these mechanistic interactions are shaped by evolution in comparative studies.

Wood frog larvae with low basal heat tolerance demonstrated a large hardening effect suggesting a trade-off between the two traits. There is an inherent link between CT_max_ and hardening which may bias the presence of the trade-off hypothesis (van Heerwaarden and Kellermann, 2020), and Deery et al. (2021) proposed that correlative evidence of the trade-off hypothesis is a statistical artifact. However, we believe our methodology of using different individuals for CT_max_ and hardening removed the risks of spurious correlation and strengthened our analyses. Based on our results, we propose that larval wood frogs support the trade-off hypothesis after a relatively short acclimation period. Hardening benefits cool-acclimated populations in response to acute heat stress but plasticity in basal heat tolerance in response to prolonged warming are likely to be more beneficial in reducing overheating risk.

## Acknowledgements

We would like to thank Jared Bilak, Abigail Pratt, Matthew Walker, and Justin Remmers for feedback and comments on our manuscript. We would like to thank the two anonymous reviewers for helpful feedback on this manuscript.

## Author contributions

Conceptualization: J.D.; Methodology: J.D.; Formal analysis: J.D.; Investigation: J.D.; Data curation: J.D.; Writing - original draft: J.D.; Writing - review & editing: J.D., R.W.; Supervision: R.W.; Project administration: J.D.; Funding acquisition: R.W.

## References

Angilletta Jr, M. J. (2009). Thermal adaptation: a theoretical and empirical synthesis: Oxford University Press.

Bahrndorff, S., Mariën, J., Loeschcke, V. and Ellers, J. (2009). Dynamics of heat-induced thermal stress resistance and Hsp 70 expression in the Springtail, Orchesella cincta. Functional Ecology, 233–239.

Barley, J. M., Cheng, B. S., Sasaki, M., Gignoux-Wolfsohn, S., Hays, C. G., Putnam, A. B., Sheth, S., Villeneuve, A. R. and Kelly, M. (2021). Limited plasticity in thermally tolerant ectotherm populations: evidence for a trade-off. Proc Biol Sci 288, 20210765.

Bellard, C., Bertelsmeier, C., Leadley, P., Thuiller, W. and Courchamp, F. (2012). Impacts of climate change on the future of biodiversity. Ecol Lett 15, 365–377.

Bettencourt, B. R., Feder, M. E. and Cavicchi, S. (1999). Experimental evolution of Hsp70 expression and thermotolerance in Drosophila melanogaster. Evolution 53, 484–492.

Blair, S. D. and Glover, C. N. (2019). Acute exposure of larval rainbow trout (Oncorhynchus mykiss) to elevated temperature limits hsp70b expression and influences future thermotolerance. Hydrobiologia 836, 155–167.

Blaustein, A. R., Walls, S. C., Bancroft, B. A., Lawler, J. J., Searle, C. L. and Gervasi, S. S. (2010). Direct and Indirect Effects of Climate Change on Amphibian Populations. Diversity 2, 281–313.

Blois, J. L., Zarnetske, P. L., Fitzpatrick, M. C. and Finnegan, S. (2013). Climate change and the past, present, and future of biotic interactions. Science 341, 499–504.

Bodensteiner, B. L., Agudelo-Cantero, G. A., Arietta, A. Z. A., Gunderson, A. R., Munoz, M. M., Refsnider, J. M. and Gangloff, E. J. (2020). Thermal adaptation revisited: How conserved are thermal traits of reptiles and amphibians? J Exp Zool A Ecol Integr Physiol.

Bowler, K. (2005). Acclimation, heat shock and hardening. Journal of Thermal Biology 30, 125–130.

Campbell Grant, E. H., Miller, D. A. W. and Muths, E. (2020). A Synthesis of Evidence of Drivers of Amphibian Declines. Herpetologica 76.

Cox, N., Young, B. E., Bowles, P., Fernandez, M., Marin, J., Rapacciuolo, G., Bohm, M., Brooks, T. M., Hedges, S. B., Hilton-Taylor, C. et al. (2022). A global reptile assessment highlights shared conservation needs of tetrapods. Nature.

Cupp Jr, P. V. (1980). Thermal tolerance of five salientian amphibians during development and metamorphosis. Herpetologica, 234–244.

Dalvi, R. S., Pal, A. K., Tiwari, L. R. and Baruah, K. (2012). Influence of acclimation temperature on the induction of heat-shock protein 70 in the catfish Horabagrus brachysoma (Gunther). Fish Physiology and Biochemistry 38, 919–927.

Deery, S. W., Rej, J. E., Haro, D. and Gunderson, A. R. (2021). Heat hardening in a pair of Anolis lizards: constraints, dynamics and ecological consequences. J Exp Biol 224.

Denny, M. W. and Dowd, W. W. (2012). Biophysics, environmental stochasticity, and the evolution of thermal safety margins in intertidal limpets. J Exp Biol 215, 934–47.

Easton, D. P., Rutledge, P. S. and Spotila, J. R. (1987). Heat shock protein induction and induced thermal tolerance are independent in adult salamanders. Journal of Experimental Zoology 241, 263–267.

Enriquez-Urzelai, U., Kearney, M. R., Nicieza, A. G. and Tingley, R. (2019). Integrating mechanistic and correlative niche models to unravel range-limiting processes in a temperate amphibian. Glob Chang Biol 25, 2633–2647.

Feder, M. E. and Hofmann, G. E. (1999). Heat-shock proteins, molecular chaperones, and the stress response: evolutionary and ecological physiology. Annual Review of Physiology 61, 243–282.

Gao, J., Zhang, W., Dang, W., Mou, Y., Gao, Y., Sun, B. J. and Du, W. G. (2014). Heat shock protein expression enhances heat tolerance of reptile embryos. Proc Biol Sci 281, 20141135.

Gilbert, A. L. and Miles, D. B. (2019). Antagonistic Responses of Exposure to Sublethal Temperatures: Adaptive Phenotypic Plasticity Coincides with a Reduction in Organismal Performance. Am Nat 194, 344–355.

Gleason, L. U. and Burton, R. S. (2015). RNA-seq reveals regional differences in transcriptome response to heat stress in the marine snail Chlorostoma funebralis. Mol Ecol 24, 610–27.

Gosner, K. L. (1960). A Simplified Table for Staging Anuran Embryos and Larvae with Notes on Identification. Herpetologica 16, 183–190.

Gu, L. L., Li, M. Z., Wang, G. R. and Liu, X. D. (2019). Multigenerational heat acclimation increases thermal tolerance and expression levels of Hsp70 and Hsp90 in the rice leaf folder larvae. J Therm Biol 81, 103–109.

Healy, T. M. and Schulte, P. M. (2012). Factors affecting plasticity in whole-organism thermal tolerance in common killifish (Fundulus heteroclitus). J Comp Physiol B 182, 49–62.

Hoekstra, L. A. and Montooth, K. L. (2013). Inducing extra copies of the Hsp70 gene in Drosophila melanogaster increases energetic demand. BMC evolutionary biology 13, 1–11.

Huey, R. B. and Stevenson, R. (1979). Integrating thermal physiology and ecology of ectotherms: a discussion of approaches. American Zoologist 19, 357–366.

Jensen, L. T., Cockerell, F. E., Kristensen, T. N., Rako, L., Loeschcke, V., McKechnie, S. W. and Hoffmann, A. A. (2010). Adult heat tolerance variation in Drosophila melanogaster is not related to Hsp70 expression. J Exp Zool A Ecol Genet Physiol 313, 35–44.

Jin, J., Zhao, M., Wang, Y., Zhou, Z., Wan, F. and Guo, J. (2019). Induced Thermotolerance and Expression of Three Key Hsp Genes (Hsp70, Hsp21, and sHsp21) and Their Roles in the High Temperature Tolerance of Agasicles hygrophila. Front Physiol 10, 1593.

Krebs, R. A. and Feder, M. E. (1998). Hsp70 and larval thermotolerance in Drosophila melanogaster: how much is enough and when is more too much? Journal of insect physiology 44, 1091–1101.

Lapwong, Y., Dejtaradol, A. and Webb, J. K. (2021a). Plasticity in thermal hardening of the invasive Asian house gecko. Evolutionary Ecology.

Lapwong, Y., Dejtaradol, A. and Webb, J. K. (2021b). Shifts in thermal tolerance of the invasive Asian house gecko (Hemidactylus frenatus) across native and introduced ranges. Biological Invasions 23, 989–996.

Li, H., Wang, Z., Mei, W. and Ji, X. (2009). Temperature acclimation affects thermal preference and tolerance in three Eremias lizards (Lacertidae). Current Zoology 55, 258–265.

Lowe, W. H., Martin, T. E., Skelly, D. K. and Woods, H. A. (2021). Metamorphosis in an Era of Increasing Climate Variability. Trends Ecol Evol 36, 360–375.

Maness, J. D. and Hutchison, V. H. (1980). Acute adjustment of thermal tolerance in vertebrate ectotherms following exposure to critical thermal maxima. Journal of Thermal Biology 5, 225–233.

Menke, M. E. and Claussen, D. L. (1982). Thermal acclimation and hardening in tadpoles of the bullfrog, Rana catesbeiana. Journal of Thermal Biology 7, 215–219.

Moyen, N. E., Crane, R. L., Somero, G. N. and Denny, M. W. (2020). A single heat-stress bout induces rapid and prolonged heat acclimation in the California mussel, Mytilus californianus. Proc Biol Sci 287, 20202561.

Phillips, B. L., Muñoz, M. M., Hatcher, A., Macdonald, S. L., Llewelyn, J., Lucy, V., Moritz, C. and Grémillet, D. (2016). Heat hardening in a tropical lizard: geographic variation explained by the predictability and variance in environmental temperatures. Functional Ecology 30, 1161–1168.

Rohr, J. R., Civitello, D. J., Cohen, J. M., Roznik, E. A., Sinervo, B. and Dell, A. I. (2018). The complex drivers of thermal acclimation and breadth in ectotherms. Ecol Lett 21, 1425–1439.

Rollins-Smith, L. A. (2017). Amphibian immunity–stress, disease, and climate change. Developmental & Comparative Immunology 66, 111–119.

Ruthsatz, K., Dausmann, K. H., Peck, M. A. and Glos, J. (2022). Thermal tolerance and acclimation capacity in the European common frog (Rana temporaria) change throughout ontogeny. J Exp Zool A Ecol Integr Physiol.

Rutledge, P., Spotila, J. and Easton, D. (1987). Heat hardening in response to two types of heat shock in the lungless salamanders Eurycea bislineata and Desmognathus ochrophaeus. Journal of Thermal Biology 12, 235–241.

Seneviratne, S., Zhang, I. X., Adnan, M., Badi, W., Dereczynski, C., Di Luca, A., Ghosh, S., Iskandar, I., Kossin, J., Lewis, S. et al. (2021). Weather and Climate Extreme Events in a Changing Climate. In: Climate Change 2021: The Physical Science Basis. In Climate Change 2021: The Physical Science Basis. Contribution of Working Group I to the Sixth Assessment Report of the Intergovernmental Panel on Climate Change eds. V. Masson-Delmotte P. Zhai A. Pirani S. L. Connors C. Péan S. Berger N. Caud Y. Chen L. Goldfarb M. I. Gomis et al.): Cambridge University Press.

Sgro, C. M., Overgaard, J., Kristensen, T. N., Mitchell, K. A., Cockerell, F. E. and Hoffmann, A. A. (2010). A comprehensive assessment of geographic variation in heat tolerance and hardening capacity in populations of Drosophila melanogaster from eastern Australia. J Evol Biol 23, 2484–93.

Sherman, E. and Levitis, D. (2003). Heat hardening as a function of developmental stage in larval and juvenile Bufo americanus and Xenopus laevis. Journal of Thermal Biology 28, 373–380.

Simon, M. N., Ribeiro, P. L. and Navas, C. A. (2015). Upper thermal tolerance plasticity in tropical amphibian species from contrasting habitats: implications for warming impact prediction. J Therm Biol 48, 36–44.

Somero, G. N. (2010). The physiology of climate change: how potentials for acclimatization and genetic adaptation will determine ‘winners’ and ‘losers’. J Exp Biol 213, 912–20.

Sørensen, J. G., Kristensen, T. N. and Loeschcke, V. (2003). The evolutionary and ecological role of heat shock proteins. Ecology Letters 6, 1025–1037.

Sørensen, J. G., Pekkonen, M., Lindgren, B., Loeschcke, V., Laurila, A. and Merilä, J. (2009). Complex patterns of geographic variation in heat tolerance and Hsp70 expression levels in the common frog Rana temporaria. Journal of Thermal Biology 34, 49–54.

Stillman, J. H. (2003). Acclimation capacity underlies susceptibility to climate change. Science 301, 65–65.

Turriago, J. L., Tejedo, M., Hoyos, J. M. and Bernal, M. H. (2022). The effect of thermal microenvironment in upper thermal tolerance plasticity in tropical tadpoles. Implications for vulnerability to climate warming. J Exp Zool A Ecol Integr Physiol 337, 746–759.

van Heerwaarden, B. and Kellermann, V. (2020). Does Plasticity Trade Off With Basal Heat Tolerance? Trends Ecol Evol 35, 874–885.

Zatsepina, O., Ulmasov, K. A., Beresten, S., Molodtsov, V., Rybtsov, S. and Evgen’Ev, M. (2000). Thermotolerant desert lizards characteristically differ in terms of heat-shock system regulation. Journal of Experimental Biology 203, 1017–1025.

Zatsepina, O. G., Velikodvorskaia, V. V., Molodtsov, V. B., Garbuz, D., Lerman, D. N., Bettencourt, B. R., Feder, M. E. and Evgenev, M. B. (2001). A Drosophila melanogaster strain from sub-equatorial Africa has exceptional thermotolerance but decreased Hsp70 expression. Journal of Experimental Biology 204, 1869–1881.

